# Human serum triglycerides promote *Staphylococcus aureus* biofilm formation and antibiotic tolerance

**DOI:** 10.64898/2026.03.05.709992

**Authors:** Elizabeth V. K. Ledger, Niamh E. Horgan, Denis Lynch, Lorraine M. Bateman, Ruth C. Massey

**Affiliations:** School of Microbiology, University College Cork, Cork, Ireland; APC Microbiome Ireland, University College Cork, Cork, Ireland; School of Chemistry, Analytical and Biological Chemistry Research Facility, University College Cork, Cork, Ireland; School of Pharmacy, University College Cork, Cork, Ireland; School of Cellular and Molecular Medicine, University of Bristol, Bristol, UK

## Abstract

Biofilm formation and antibiotic tolerance are major contributors to the persistence of *Staphylococcus aureus* infections, yet how the host environment affects these phenotypes remains poorly understood. Here, we show that incubation in human serum primes *S. aureus* to form robust biofilms and tolerate vancomycin and daptomycin, last resort antibiotics for the treatment of antibiotic-resistant staphylococcal infections. Mechanistically, we demonstrate that the staphylococcal Geh lipase is essential for serum-induced biofilm formation by liberating glycerol from host lipids, which is then used to promote increased synthesis of D-alanylated wall teichoic acids, driving biofilm development. Inhibition of the Geh lipase or wall teichoic acid synthesis markedly reduces biofilm formation and restores antibiotic susceptibility, highlighting clinically achievable strategies to inhibit host-induced biofilm formation and prevent the associated antibiotic tolerance. Together, our findings reveal a host-driven mechanism of biofilm-associated antibiotic tolerance in *S. aureus* and provide rational targets for therapeutic intervention.

## Introduction

*Staphylococcus aureus* is a major human pathogen that causes a range of infections from minor skin and soft tissue infections to invasive diseases like bacteraemia, which is associated with a 30% mortality rate and approximately 300,000 deaths annually^1^. One of the main reasons for this high mortality is the propensity of the bacterium to disseminate from the bloodstream and cause secondary infections, including endocarditis, osteomyelitis and deep tissue abscesses^1^. These infections frequently have a biofilm component, where multicellular communities are embedded within a self-produced matrix of proteins, polysaccharides, DNA and lipids^2^. Biofilms reduce the efficacy of antibiotics and protect the bacteria within them from the immune system, making biofilm-based infections very difficult to eradicate. As a result, patients often suffer from chronic or relapsing infections, significantly contributing to morbidity and mortality.

Understanding how these biofilms form is crucial for developing new approaches to treat these infections more effectively. However, most current knowledge of biofilm formation comes from research on bacteria grown in laboratory media, meaning that the host factors that influence biofilm formation are poorly understood. In particular, very little is known about how exposure to the bloodstream environment, which *S. aureus* must transit through before reaching deeper tissues, influences the bacterium’s subsequent ability to form biofilms at secondary sites.

Many strains of *S. aureus* are poor biofilm formers *in vitro*, and glucose or NaCl must be added to media to induce biofilm formation. Under these conditions, many cellular components contribute to the biofilm formation, including the wall teichoic acids (WTA), surface polymers important for attachment to surfaces, surface proteins, including many sortase-anchored LPxTG proteins, and polysaccharide intercellular adhesin (PIA). However, the conditions found in these *in vitro* assays do not accurately replicate the conditions found during infection, in part due to the lack of host-derived nutrients and macromolecules. One structure which is profoundly impacted by the host environment is the cell envelope, which as the primary interface between *S. aureus* and host cells and surfaces, is a major determinant of biofilm formation.

The host environment affects both the structure of the cell wall and the composition and properties of the cell membrane. For example, *S. aureus* incorporates serum-derived fatty acids into its phospholipids^3,4^, altering membrane fluidity^5,6^, and becomes coated in a layer of host proteins and lipids^4,7^. In addition, the bloodstream environment triggers extensive cell wall remodelling, consisting of a significant accumulation of both peptidoglycan and WTA, and leading to *in vivo*-grown bacteria having much thicker cell walls than laboratory media-grown cells^8,9^. This cell wall thickening is due to host antimicrobial peptides activating the GraRS two-component signalling system, reducing peptidoglycan hydrolysis, and requires the penicillin-binding protein PBP4^8,10^. The thickened cell wall has profound effects on bacterial survival, as it protects the bacteria from membrane-targeting antibiotics including daptomycin and conceals many surface proteins, protecting the bacteria from opsonophagocytosis by neutrophils^8,11^.

However, despite the central role of these host-induced cell envelope changes in bacterial survival in the bloodstream, their consequences for biofilm formation remain unknown. As these adaptations require exposure to host conditions and develop over time, they are absent from conventional *in vitro* biofilm assays, meaning that we do not have the full picture of how biofilms form under host conditions.

Here, using human serum to model the bloodstream environment, we show that host conditions prime *S. aureus* to produce more biofilm and show increased antibiotic tolerance. We uncover the mechanism behind this, identifying bacterial metabolism of host lipids as a crucial biofilm-promoting process. These findings provide important insights into how biofilms may form *in vivo* and identify a potential new therapeutic combination approach to reduce biofilm formation and improve antibiotic efficacy.

## Results

### Serum triggers increased biofilm formation and antibiotic tolerance

During *S. aureus* bacteraemia, bacteria frequently leave the bloodstream and cause infections at secondary sites, including deep tissue abscesses, osteomyelitis and infective endocarditis^1^. These infections are often biofilm based, protecting the bacteria within from the immune system and antibiotics and making them difficult to eradicate. Biofilm formation is frequently studied using bacteria which have been grown in laboratory media^12^. However, this does not replicate the *in vivo* environment, where bacteria have had time to adapt to the bloodstream environment entering tissues and forming biofilms. Therefore, to investigate the effect that this adaptation to the host environment has on biofilm formation, we studied the biofilm formation of *S. aureus* in two states: i) grown in tryptic soy broth (TSB) to mimic *in vitro* conditions commonly used experimentally (“TSB-grown”) and ii) incubated in normal human serum for 16 h to mimic *in vivo* conditions (“serum-incubated”).

The USA300 JE2 laboratory strain of *S. aureus* was grown to either mid-exponential phase or stationary phase in TSB before being incubated, or not, in human serum for 16 h. After this serum incubation, cultures were washed to remove serum and then inoculated into TSB supplemented with glucose and incubated for 24 h to enable biofilm formation before quantification of biofilm formation by crystal violet staining. The CFU count did not increase during the incubation in serum and so equal numbers of CFUs were inoculated in each case (Fig. S1). *S. aureus* formed low levels of biofilm when grown in TSB (A_595_ < 0.5), whereas pre-incubation in serum led to a significant increase in subsequent biofilm biomass regardless of growth phase (Fig. 1a). As no difference was observed between the growth phases, only exponential phase cultures were used for the remainder of this work. This serum-induced biofilm formation was also observed in three other commonly used laboratory strains, SH1000, Col and Newman, although the ability of serum to trigger biofilm formation in Newman was lower than the other strains (Fig. 1b). It was also observed in all five tested clinical endocarditis isolates^13^, comprising both MRSA and MSSA strains from clonal complexes 22 and 30, representative of the most commonly circulating strains in the UK and Ireland (Fig. 1c).

**Figure 1.**
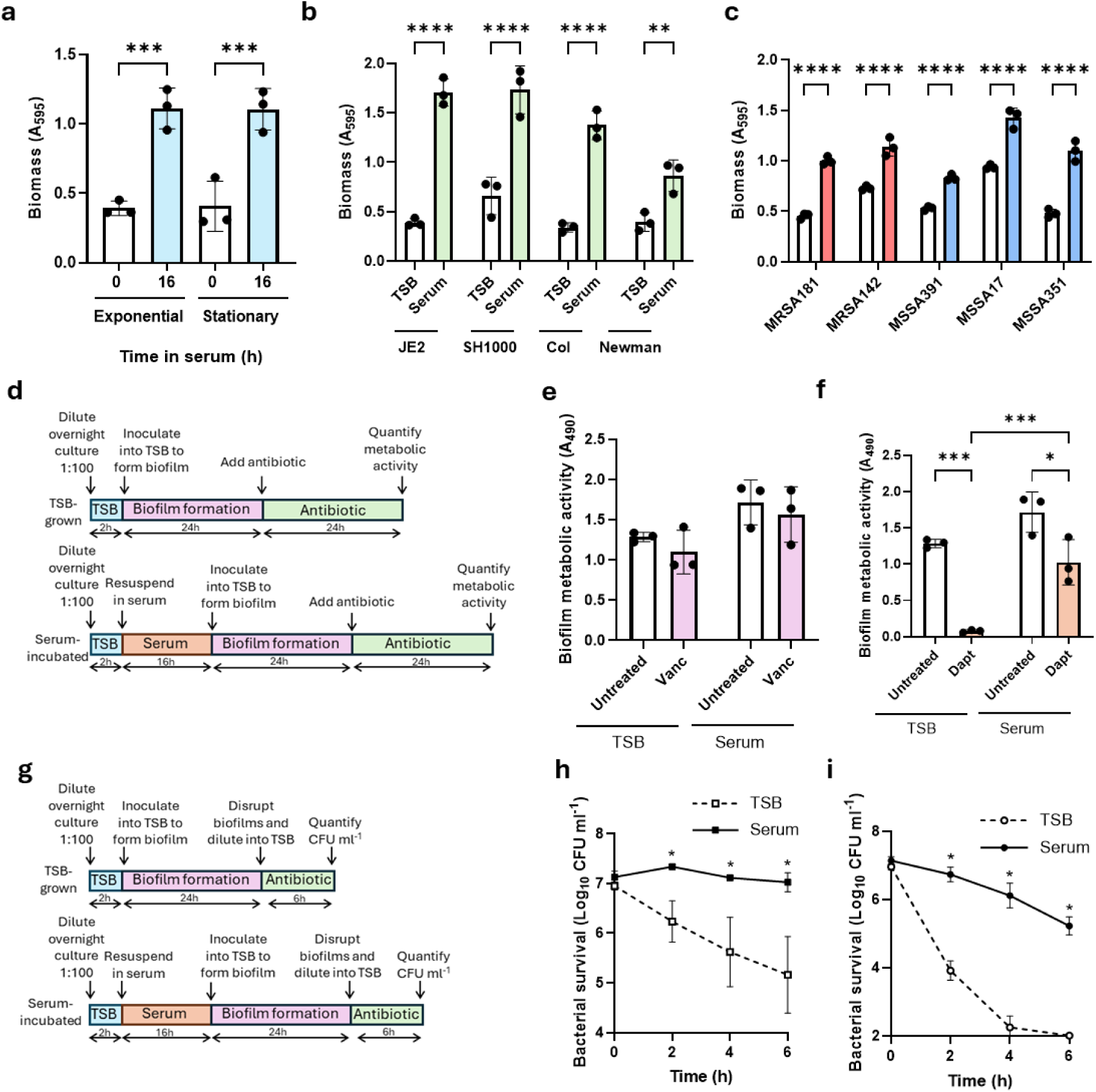
Serum triggers increased biofilm formation and antibiotic tolerance. (a) Biomass of biofilms formed from *S. aureus* grown to mid-exponential phase or stationary phase and then incubated, or not, for 16 h in human serum, as determined by crystal violet staining. (b) Biomass of biofilms formed from four different laboratory strains of *S. aureus* after being TSB-grown or serum-incubated as measured by crystal violet staining. (c) Biomass of biofilms formed from five different clinical bacteraemia isolates of *S. aureus* after being TSB-grown or serum-incubated as measured by crystal violet staining. (d) Schematic outlining the protocol used to investigate the susceptibility of intact biofilms to antibiotics. Metabolic activity of biofilms formed from TSB-grown and serum-incubated JE2 after a 24 h exposure to (e) 20 µg ml^-1^ vancomycin or (f) 20 µg ml^-1^ daptomycin. (g) Schematic outlining the protocol used to investigate the antibiotic susceptibility of disrupted biofilms. Bacterial survival as measured by log_10_ CFU ml^-1^ of JE2 isolated from disrupted biofilms formed from TSB-grown or serum-incubated cells after a 6 h exposure to (h) 20 µg ml^-1^ vancomycin or (i) 20 µg ml^-1^ daptomycin. Data in a, b, c, e and f represent the mean ± standard deviation of three independent repeats. Data in h and i represent the geometric mean ± geometric standard deviation of three independent repeats. Data in a, b, c, h and i were analysed by two-way ANOVA with Sidak’s *post-hoc* test. Data in e and f were analysed by two-way ANOVA with Tukey’s *post-hoc* test. (*, P < 0.05, **, P < 0.01, ***, P < 0.001 and ****, P < 0.0001).

Staphylococcal biofilms have previously been reported to be predominantly protein based^14^ and so to test whether this was also the case for biofilms generated from serum-incubated cells we treated formed biofilms with proteinase K, DNase or sodium metaperiodate to degrade proteins, DNA and carbohydrates, respectively, before quantifying biomass. DNase and sodium metaperiodate had no effect on the biomass of biofilms formed from either TSB-grown or serum-incubated cells (Fig. S2). By contrast, protease treatment led to a significant loss of biomass, indicating that the biofilms formed by serum-incubated cells are also mainly composed of proteins (Fig. S2).

We next investigated whether the biofilms formed from serum-incubated *S. aureus* showed altered antibiotic tolerance compared to those formed from TSB-grown cells. To do this, TSB-grown and serum-incubated biofilms were generated as above, before the supernatant was removed and replaced with fresh TSB or TSB supplemented with 20 µg ml^-1^ vancomycin or daptomycin. Biofilms were incubated with antibiotics for 24 h before the metabolic activity of the biofilms was determined with XTT (Fig. 1d). Vancomycin showed no activity against either TSB-grown or serum-incubated biofilms, likely as it is unable to penetrate the biofilm (Fig. 1e). Daptomycin was highly active against TSB-grown biofilms, with negligible metabolic activity observed after the 24 h treatment (Fig. 1f). By contrast, it was significantly less effective against serum-incubated biofilms, where high levels of metabolic activity remained after antibiotic exposure (Fig. 1f).

Finally, we investigated whether the bacteria within the biofilms showed altered antibiotic susceptibility by mechanically disrupting TSB-grown and serum-incubated biofilms, exposing the bacteria to antibiotics and measuring survival by CFU counts (Fig. 1g). In the disrupted biofilms, TSB-grown bacteria were susceptible to both vancomycin and daptomycin, with 2-logs and 5-logs of killing observed, respectively, by 6 h. By contrast, incubation in serum protected *S. aureus* from both antibiotics, with no killing observed by vancomycin and less than 2-logs of killing observed by daptomycin (Fig. 1h – i).

Taken together, incubation in human serum primes *S. aureus* to produce more biofilm and enhances antibiotic tolerance, both within intact biofilms and in the bacteria that are released from them, suggesting that host adaptation may play a key role in the persistence and treatment resistance of invasive infections.

### Lipase-mediated host lipid metabolism is required for biofilm formation

The next objective was to understand why incubation in serum led to increased biofilm formation. The first hypothesis was that host factors from serum were binding to the bacterial surface and enhancing bacterial adhesion and biofilm formation. To test this, *S. aureus* was incubated in serum for 30 min to enable host factors to bind before biofilm formation was measured. This 30 min serum incubation led to a very slight but non-significant increase in biofilm formation, and biomass was significantly lower than that of 16 h serum-incubated cells, suggesting that the binding of host factors was not sufficient to explain the increased biofilm formation (Fig. S3).

*S. aureus* uses proteinaceous and non-proteinaceous cell wall macromolecules to attach to surfaces and form biofilms. Many cell wall-anchored proteins produced by *S. aureus* have been demonstrated to contribute to surface attachment^15^. As many of these proteins are attached to the cell wall by sortase A, we investigated the ability of a *srtA*::Tn mutant to form biofilm after serum incubation. The *srtA*::Tn mutant formed similar levels of biofilm to the WT strain, indicating that cell wall anchored proteins are not required for biofilm formation under these conditions (Fig. S4). The *icaABCD* operon, which encodes the proteins responsible for synthesising the polysaccharide PIA has also been shown to be important for biofilm condition under some conditions^16^, however, it was not required for serum-induced biofilm formation (Fig. S5).

To understand which serum component was responsible for triggering increased biofilm formation, we subjected serum to fractionation through a 10 kDa membrane, heat-inactivation to remove heat-labile proteins and delipidation to remove lipids, before measuring biofilm formation. Both the <10 kDa fraction and delipidated serum (DLS) were unable to trigger increased biofilm formation, demonstrating that the component responsible was >10 kDa and was lipid based (Fig. 2a). In serum, most lipids are found bound to host proteins, especially albumin and lipoproteins, possibly explaining why the biofilm-inducing activity was retained in the >10 kDa fraction despite lipids themselves being much smaller than this.

**Figure 2.**
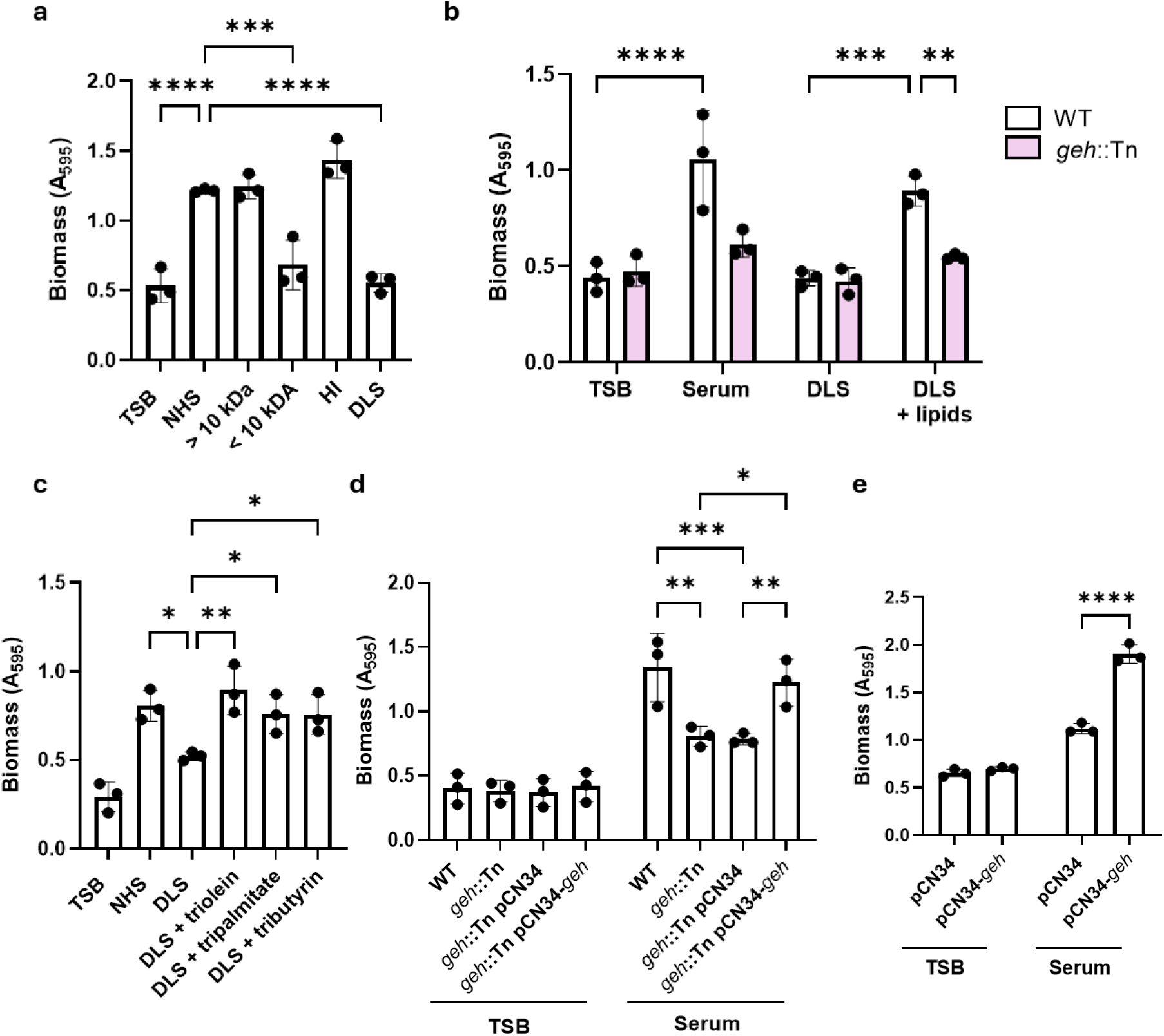
Lipase-mediated host lipid metabolism is required for biofilm formation. (a) Biomass of biofilms formed from *S. aureus* grown in TSB or incubated for 16 h in normal human serum (NHS), serum fractionated through a 10 kDa filter, heat-inactivated (HI) serum or delipidated serum (DLS) as determined by crystal violet staining. (b) Biomass of biofilms formed from JE2 WT or the *geh*::Tn mutant after being TSB-grown or incubated for 16 h in serum, DLS or DLS supplemented with serum-extracted lipids as measured by crystal violet staining. (c) Biomass of biofilms formed from JE2 WT after being TSB-grown or incubated for 16 h in NHS, DLS or DLS supplemented with 1 mM triolein, tripalmitate or tributyrin as measured by crystal violet staining. (d) Biomass of biofilms formed from TSB-grown and serum-incubated cultures of JE2 WT, the *geh*::Tn mutant, the *geh*::Tn mutant carrying the empty pCN34 plasmid or the *geh*::Tn mutant complemented with pCN34-*geh* as measured by crystal violet staining. (e) Biomass of biofilms formed from TSB-grown or serum-incubated cultures of Newman either carrying the empty pCN34 plasmid or complemented with pCN34-*geh* as measured by crystal violet staining. Data in all panels represent the mean ± standard deviation of three independent repeats. Data in a and c were analysed by one-way ANOVA with Dunnett’s *post-hoc* test. Data in b and e were analysed by two-way ANOVA with Sidak’s *post-hoc* test and data in d were analysed by two-way ANOVA with Tukey’s *post-hoc* test. *, P < 0.05, **, P < 0.01, ***, P < 0.001 and ****, P < 0.0001.

This suggested that lipids were the serum factor that led to increased biofilm formation. Since *S. aureus* secretes lipases that break down host lipids into fatty acids that can be incorporated into bacterial lipids by fatty acid kinase FakA/B1/B2^3^, we next investigated whether this activity was required for biofilm induction in serum, using mutants from the Nebraska Transposon Mutant Library (NTML). This showed that Geh, the predominant staphylococcal lipase, was required for biofilm formation in serum but that fatty acid kinase was not (Fig. 2b and Fig. S6). The *geh*::Tn mutant showed significantly reduced biofilm formation in normal human serum (NHS) compared to the WT, while neither strain formed high levels of biofilm in delipidated serum (DLS) (Fig. 2b). Critically, addition of lipids extracted from NHS using a Bligh-Dyer extraction to DLS was able to restore biofilm formation in the WT strain but not in the *geh*::Tn mutant (Fig. 2b). The ability of lipids to trigger biofilm formation was also demonstrated by the finding that supplementation of DLS with purified triglycerides also enhanced biofilm formation (Fig. 2c). Complementation of the *geh*::Tn mutant by the expression of *geh* under the control of its native promoter restored biofilm formation to WT levels (Fig. 2d).

Interestingly, the Newman strain tested in Fig. 1b contains a prophage inserted into the *geh* lipase gene, disrupting its expression^17^. To test whether this explained its lower level of biofilm formation compared to the other laboratory strains tested, *geh* was complemented in the Newman background, leading to a significant increase in serum-induced biofilm formation in this background as well (Fig. 2e). Together, this demonstrates that the metabolism of host lipids by the staphylococcal Geh lipase leads to enhanced biofilm formation.

### Lipase-mediated generation of glycerol promotes biofilm formation by enabling increased synthesis of D-alanylated WTA

The next objective was to determine how Geh lipase was contributing to biofilm formation in serum. Since triglycerides, which are broken down into fatty acids and glycerol by lipase, were demonstrated to enhance biofilm formation in Fig. 2c, we next added either fatty acids or glycerol into DLS to determine which led to increased biofilm formation. Five fatty acids were added individually (5 mM each) or in combination (1 mM each, total concentration 5 mM). As some fatty acids have previously been reported to show anti-bacterial activity, we first tested whether this incubation in DLS with fatty acids affected bacterial survival to ensure that equal numbers of CFU were inoculated when testing biofilm formation (Fig. S7a). Fatty acid supplementation did not affect bacterial survival, except for linoleic acid (Fig. S7a). Addition of fatty acids individually or in combination did not lead to an increase in biofilm formation (Fig. S7b and Fig. 3a). By contrast, supplementation of DLS with glycerol restored biofilm formation to the level triggered by NHS, demonstrating that it is the release of glycerol by lipase that is important for biofilm formation rather than the fatty acids (Fig. 3a). Interestingly, supplementation of TSB with glycerol did not lead to increased biofilm formation, indicating that additional factors present in the host environment are also required (Fig. S8).

**Figure 3.**
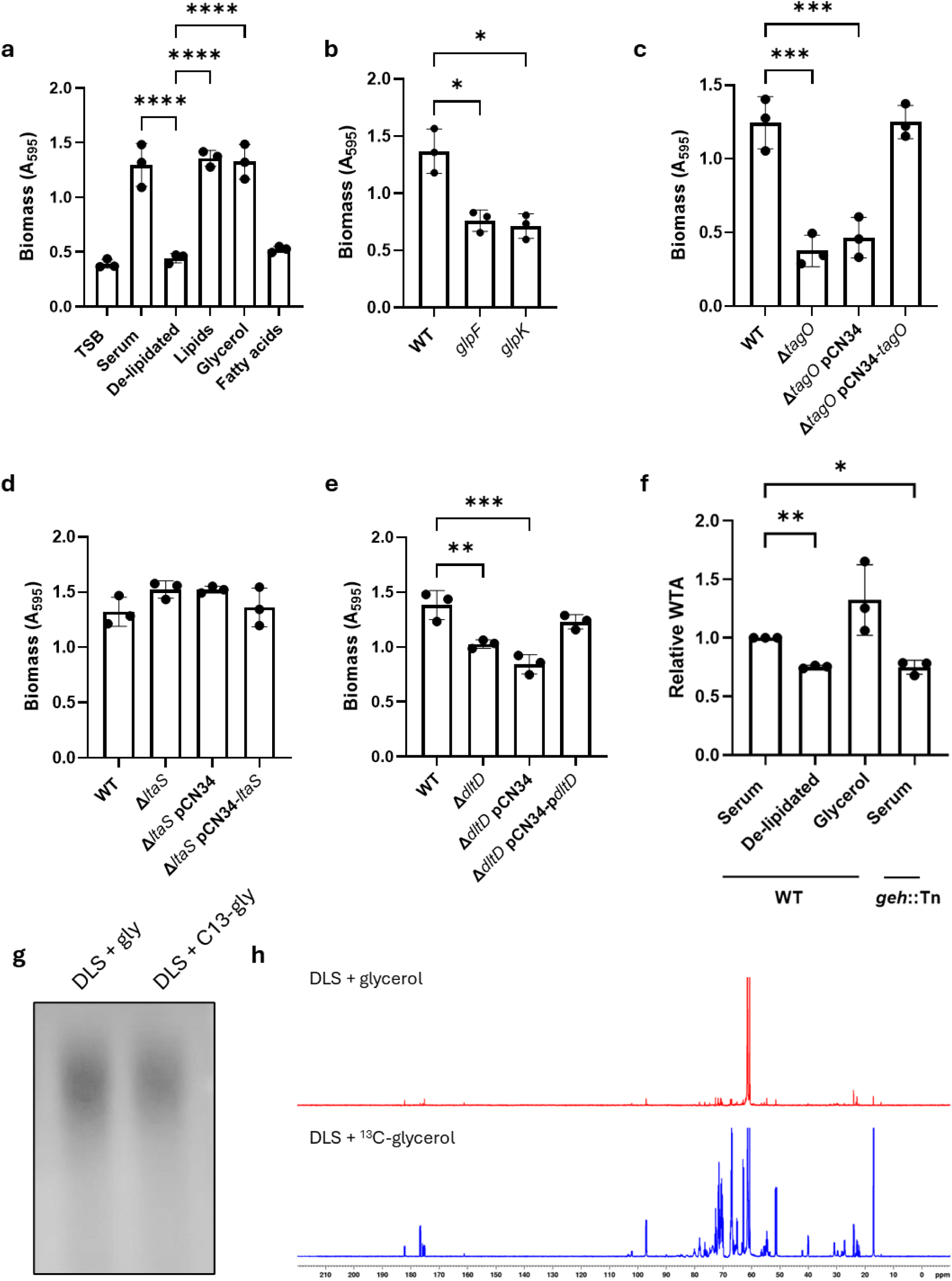
Lipase-mediated generation of glycerol promotes biofilm formation by enabling increased synthesis of D-alanylated WTA. (a) Biomass of biofilms formed from JE2 WT *g*rown in TSB or incubated for 16 h in serum or delipidated serum supplemented with serum-extracted lipids, 0.1% glycerol or 5 mM fatty acids, as determined by crystal violet staining. (b) Biomass of biofilms formed from JE2 WT or the *glpF*::Tn mutant or the *glpK*::Tn mutant after being incubated for 16 h in serum as measured by crystal violet staining. (c) Biomass of biofilms formed from serum-incubated cultures of LAC* WT, the Δ*tagO* mutant, the Δ*tagO* mutant carrying the empty pCN34 plasmid or the Δ*tagO* mutant complemented with *pCN34-tagO* as measured by crystal violet staining. (d) Biomass of biofilms formed from serum-incubated cultures of LAC* WT, the Δ*ltaS*-S2 mutant, the Δ*ltaS*-S2 mutant carrying the empty pCN34 plasmid or the Δ*ltaS*-S2 mutant complemented with *pCN34-ltaS* as measured by crystal violet staining. (e) Biomass of biofilms formed from serum-incubated cultures of LAC* WT, the Δ*dltD* mutant, the Δ*dltD* mutant carrying the empty pCN34 plasmid or the Δ*dltD* mutant complemented with *pCN34-dltD* as measured by crystal violet staining. (f) Quantification of WTA extracted from JE2 WT or the *geh*::Tn mutant after growth in serum or delipidated serum supplemented or not with 0.1% glycerol. (g) Native PAGE analysis of WTA extracted from JE2 WT after incubation in delipidated serum supplemented with 0.1% glycerol or 0.1% ^13^C-glycerol. (h) NMR spectra of WTA extracts from JE2 WT after incubation in delipidated serum supplemented with 0.1% glycerol or 0.1% ^13^C-glycerol. The signals at 60.7 and 61.4 ppm are from tris in the extraction buffer; the spectra are calibrated to the tris CH2 signal at 61.4 ppm and both spectra are scaled to this signal (exactly overlaying spectra are shown in Fig. S14). Data in graphs represent the mean ± standard deviation of three independent repeats. Spectra in h are representative of two independent repeats. Data in a, b, c, d, e and f were analysed by one-way ANOVA with Dunnett’s *post-hoc* test. *, P < 0.05, **, P < 0.01, ***, P < 0.001 and ****, P < 0.0001.

To be used by *S. aureus,* glycerol must be imported and phosphorylated to generate glycerol-6-phosphate, processes performed by glycerol uptake facilitator (GlpF) and glycerol kinase (GlpK), respectively. Therefore, to gain additional confidence that glycerol was contributing to biofilm formation in serum, we tested the biofilm formation of serum-incubated *glpF*::Tn and *glpK*::Tn mutants. Serum triggered significantly less biofilm formation in each mutant compared to the WT strain (Fig. 3b), confirming that glycerol import and phosphorylation are required for serum-induced biofilm formation.

We next aimed to determine why glycerol enhanced biofilm formation, hypothesising that *S. aureus* could be using it as a precursor for a key cell envelope component that enhances biofilm formation. *S. aureus* is known to use glycerol to make its phospholipids^18^, however, it also synthesises two glycerol-containing surface polymers, wall teichoic acids (WTA) and lipoteichoic acids (LTA)^19,20^. In particular, WTA has been found to be important for the initial attachment of bacteria to surfaces, the first step of biofilm formation^21^. We found that WTA is crucial for biofilm formation in serum, with a mutant defective for WTA synthesis (Δ*tagO*) showing significantly reduced biofilm formation in serum, and complementation restoring biofilm formation to WT levels (Fig. 3c). LTA was not required for biofilm formation, with a mutant defective for LTA synthesis (Δ*ltaS*) not showing significantly reduced biofilm formation in serum compared to the WT (Fig. 3d). Teichoic acids are glycosylated by TarM and TarS and D-alanylated by DltABCD, which lowers the net negative charge of the surface and may impact binding to surfaces and biofilm formation. WTA glycosylation was not required for biofilm formation (Fig. S9), while D-alanylation did contribute, with a Δ*dltD* mutant which lacks this modification showing significantly reduced biofilm formation in serum, albeit not to the same degree as a WTA-deficient mutant (Fig. 3e).

Due to the requirement of WTA for biofilm formation in serum, we hypothesised that the glycerol released by the Geh lipase was being imported and used to synthesize WTA. To test this, we quantified WTA levels in *S. aureus* incubated in NHS, DLS and DLS supplemented with glycerol. Incubation in DLS led to significantly less WTA content than incubation in NHS in the WT strain, while the addition of glycerol into DLS led to increased WTA (Fig. 3f and Fig. S10). In agreement with the requirement of Geh for biofilm formation in serum, the *geh*::Tn mutant also had less WTA in NHS than the WT strain (Fig. 3f and Fig. S10).

Finally, we tested whether exogenously-supplied glycerol was used to make WTA by growing *S. aureus* in DLS supplemented with glycerol or with ^13^C-labelled glycerol, followed by WTA extraction (Fig. 3g) and analysis by NMR spectroscopy (Fig. 3h and Fig. S11-14). Although the extraction from the cells grown with labelled glycerol yielded slightly less WTA than that from cells grown with unlabelled glycerol (Fig. 3g), NMR analysis revealed that there was significant incorporation of carbon atoms from the ^13^C-labelled glycerol into the WTA. The presence of multiple signals showing fine splitting (^13^C-^13^C scalar coupling) indicates that the glycerol has been metabolised and incorporated into various moieties of the WTA molecules. Based on comparison with literature NMR analysis for closely analogous compounds in deuterium oxide, this incorporation likely includes glycerol phosphate (δ_C_ = 62.9 and 72.7), the D-alanine residues (δ_C_ = 17.0, 51.4 and 176.7) and the β*-*GlcNAc modification (δ_C_ = 24.0, 54.6, 97.0 and 175.2) (Fig. 3h and Table 1). As determined by integration, for these signals, there was between a ∼5 – 50-fold enrichment of ^13^C observed in the labelled WTA extraction compared to the unlabelled extraction (Table 1). Therefore, it is likely that the glycerol released by the lipase enzyme is used to support WTA synthesis. Taken together, the release of glycerol from host lipids by Geh lipase enables *S. aureus* to synthesise high levels of WTA, enhancing biofilm formation.

**Table 1.** Assignment of specific signals within the ^13^C NMR spectra based on closely related compounds reported in the literature. *^a^* Chemical shift values referenced to tris *C*H_2_ signal at 61.4 ppm^25^. *^b^* Integration values determined relative to the integration value of tris signal at 60.7 ppm. *^c^* Proportional increase in integration value calculated from ratio of integration of corresponding signals in ^13^C NMR spectra of DLS + ^13^C-glycerol sample/ DLS + glycerol sample.

| WTA component | $\delta_c$ (ppm)<sup>a</sup> | DLS + gly integration value<sup>b</sup> | DLS + $^{13}\text{C}$-gly integration value<sup>b</sup> | Relative increase in Integration | Literature $\delta_c$ value used for component signal identification<sup>c</sup> |
| --- | --- | --- | --- | --- | --- |
| D-alanyl ester |  |  |  |  |  |
| CH<sub>3</sub> | 17.0 | 0.70 | 32.55 | 46.50 | 16.5 (Alanine, in D<sub>2</sub>O)<sup>22</sup> |
| CH | 51.4 | 0.69 | 24.79 | 35.93 | 50.6 (Alanine, in D<sub>2</sub>O)<sup>22</sup> |
| C=O | 176.7 | 0.14 | 4.95 | 35.36 | 176.6 (Alanine, in D<sub>2</sub>O)<sup>22</sup> |
| Glycerol phosphate |  |  |  |  |  |
| C-1 & C-3 | 62.9 | 1.13 | 31.43 | 27.81 | 62.0 (Glycerol Cyclic Phosphate, in D<sub>2</sub>O)<sup>23</sup> |
| C-2 | 72.7 | 0.77 | 25.16 | 32.68 | 72.0 (Glycerol Cyclic Phosphate, in D<sub>2</sub>O)<sup>23</sup> |
| β-GlcNAc | | | | | |
| CH<sub>3</sub> | 24.0 | 0.98 | 4.55 | 4.64 | 22.9 (β-GlcNAc, in D<sub>2</sub>O)<sup>24</sup> |
| C-2 | 54.6 | 1.15 | 22.26 | 19.36 | 57.5 (β-GlcNAc, in D<sub>2</sub>O)<sup>24</sup> |
| C-1 | 97.0 | 1.02 | 18.40 | 18.04 | 95.7 (β-GlcNAc, in D<sub>2</sub>O)<sup>24</sup> |
| C=O | 175.2 | 0.55 | 3.01 | 5.47 | 175.5 (β-GlcNAc, in D<sub>2</sub>O)<sup>24</sup> |

### Inhibition of lipase or WTA synthesis prevents serum-induced biofilm formation and antibiotic tolerance

As a final step, we aimed to determine whether serum-induced biofilm formation and its associated antibiotic tolerance could be reduced therapeutically. To do this, we inhibited the Geh lipase with the clinically-approved anti-obesity drug orlistat and inhibited WTA synthesis with the TagO inhibitor tunicamycin.

Supplementation of serum with orlistat significantly reduced subsequent biofilm formation (Fig. 4a), further confirming that lipase activity is required for serum-induced biofilm formation and showing that this process can be inhibited therapeutically. Moreover, inhibition of lipase genetically using the *geh*::Tn mutant or via the addition of orlistat into serum increased the susceptibility of both intact and mechanically disrupted biofilms to daptomycin (Fig. 4b – c).

**Figure 4.**
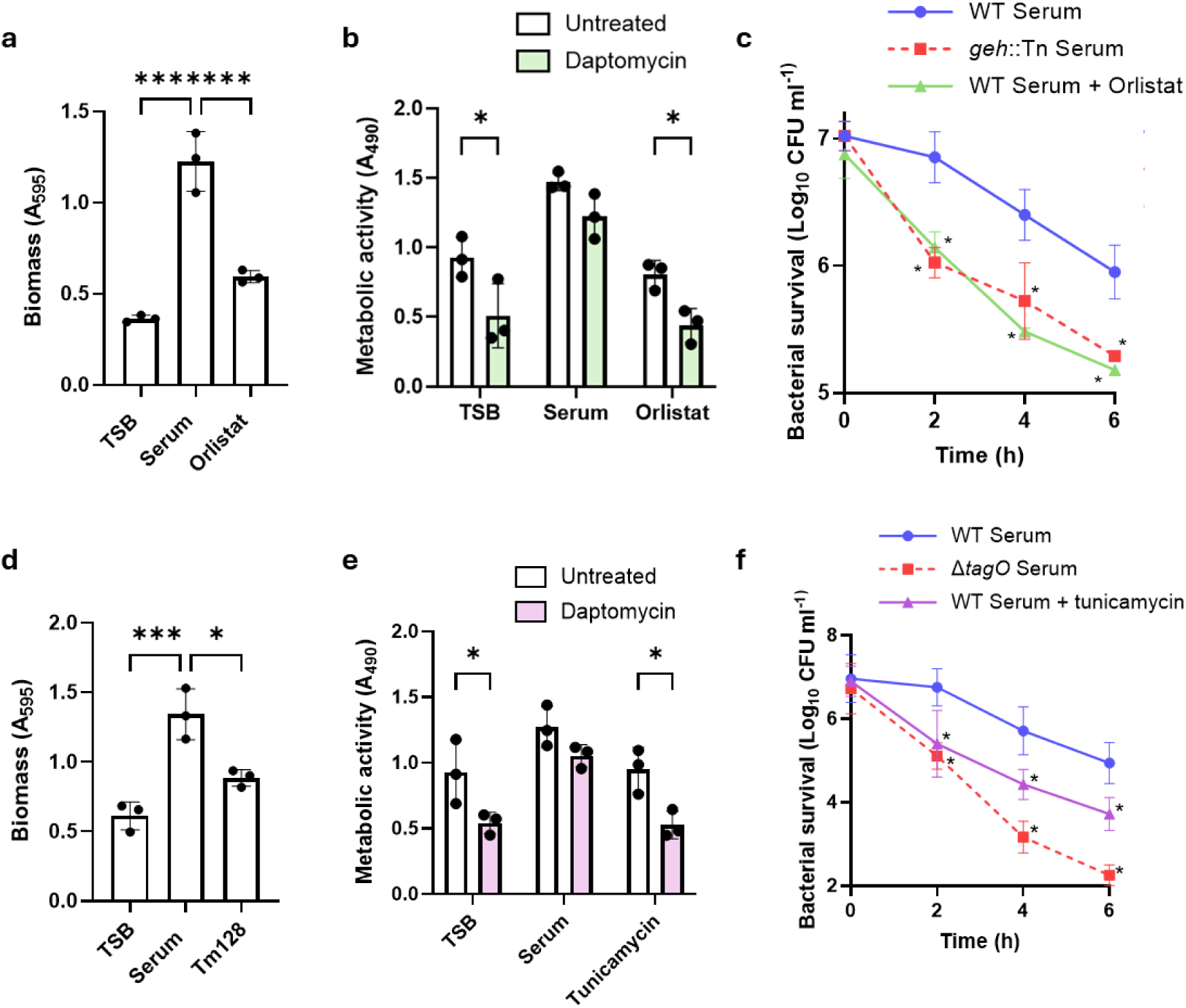
Inhibition of lipase or WTA synthesis prevents serum-induced biofilm formation and antibiotic tolerance. (a) Biomass of biofilms formed from JE2 WT grown in TSB or incubated for 16 h in serum supplemented or not with 100 µg ml^-1^ orlistat as determined by crystal violet staining. (b) Metabolic activity of biofilms formed from JE2 WT after being TSB-grown or incubated for 16 h in serum supplemented or not with 100 µg ml^-1^ orlistat and then exposed, or not, to 20 µg ml^-1^ daptomycin for 24 h. (c) Biofilms formed from JE2 WT or the *geh*::Tn mutant which had been incubated in serum supplemented or not with 100 µg ml^-1^ orlistat were disrupted and exposed to 20 µg ml^-1^ daptomycin for 6 h and log_10_ CFU ml^-1^ determined every 2 h. (d) Biomass of biofilms formed from JE2 WT *g*rown in TSB or incubated for 16 h in serum supplemented or not with 128 µg ml^-1^ tunicamycin as determined by crystal violet staining. (e) Metabolic activity of biofilms formed from JE2 WT after being TSB-grown or incubated for 16 h in serum supplemented or not with 128 µg ml^-1^ tunicamcyin and then exposed, or not, to 20 µg ml^-1^ daptomycin for 24 h. (f) Biofilms formed from LAC* WT or the Δ*tagO* mutant which had been incubated in serum supplemented or not with 128 µg ml^-1^ tunicamycin were disrupted and exposed to 20 µg ml^-1^ daptomycin for 6 h and log_10_ CFU ml^-1^ determined every 2 h. Data in panels a, b, d and e represent the mean ± standard deviation of three independent repeats and data in panels c and f represent the geometric mean ± geometric standard deviation. Data in a and d were analysed by one-way ANOVA with Dunnett’s *post-hoc* test. Data in b and e were analysed by two-way ANOVA with Sidak’s *post-hoc* test and c and f were analysed by two-way ANOVA with Dunnett’s *post-hoc* test. *, P < 0.05, **, P < 0.01, ***, P < 0.001 and ****, P < 0.0001.

We next examined whether inhibition of teichoic acid synthesis would also be a viable strategy to reduce serum-induced biofilm formation and antibiotic tolerance. Supplementation of serum with a sub-lethal concentration of tunicamycin significantly reduced biofilm formation (Fig. 4d), further confirming the importance of WTA synthesis for this process. Inhibition of WTA synthesis with tunicamycin also increased daptomycin susceptibility of intact biofilms (Fig. 4e). Furthermore, both genetic and chemical inhibition of WTA synthesis reduced the daptomycin tolerance of cells released from serum-incubated biofilms by mechanical disruption compared to the WT strain (Fig. 4f).

Taken together, serum triggers biofilm formation and antibiotic tolerance in *S. aureus* via the Geh lipase which releases glycerol from host lipids to drive enhanced D-alanylated WTA synthesis. Targeting either the lipase or WTA synthesis reduces biofilm formation and restores antibiotic susceptibility, highlighting lipase and WTA synthesis as viable targets to inhibit biofilm formation and restore antibiotic efficacy during invasive infection.

## Discussion

Treatment of invasive staphylococcal infections is challenging and is further complicated by frequent biofilm formation *in vivo* and the antibiotic recalcitrance of these biofilms^26^. Understanding how biofilms form *in vivo* is crucial to understanding the reasons behind this failure and improving the efficacy of antibiotic therapy. Here, we demonstrate that exposure to the bloodstream environment is not only a transient stage enabling dissemination but also actively primes *S. aureus* for enhanced biofilm formation and antibiotic tolerance. Using human serum to model the bloodstream, we demonstrated that adaptation to this environment induced *S. aureus* to produce high amounts of biofilm which was highly tolerant to the last resort antibiotic daptomycin. Mechanistically, this is due to the staphylococcal Geh lipase liberating glycerol from host triglycerides, which is used to synthesise increased levels of WTA, enhancing biofilm formation (Fig. 5). This links the metabolism of host lipids directly to cell envelope remodelling and biofilm formation and raises the possibility that inhibition of lipid metabolism may be a viable strategy to reduce biofilm formation.

**Figure 5.**
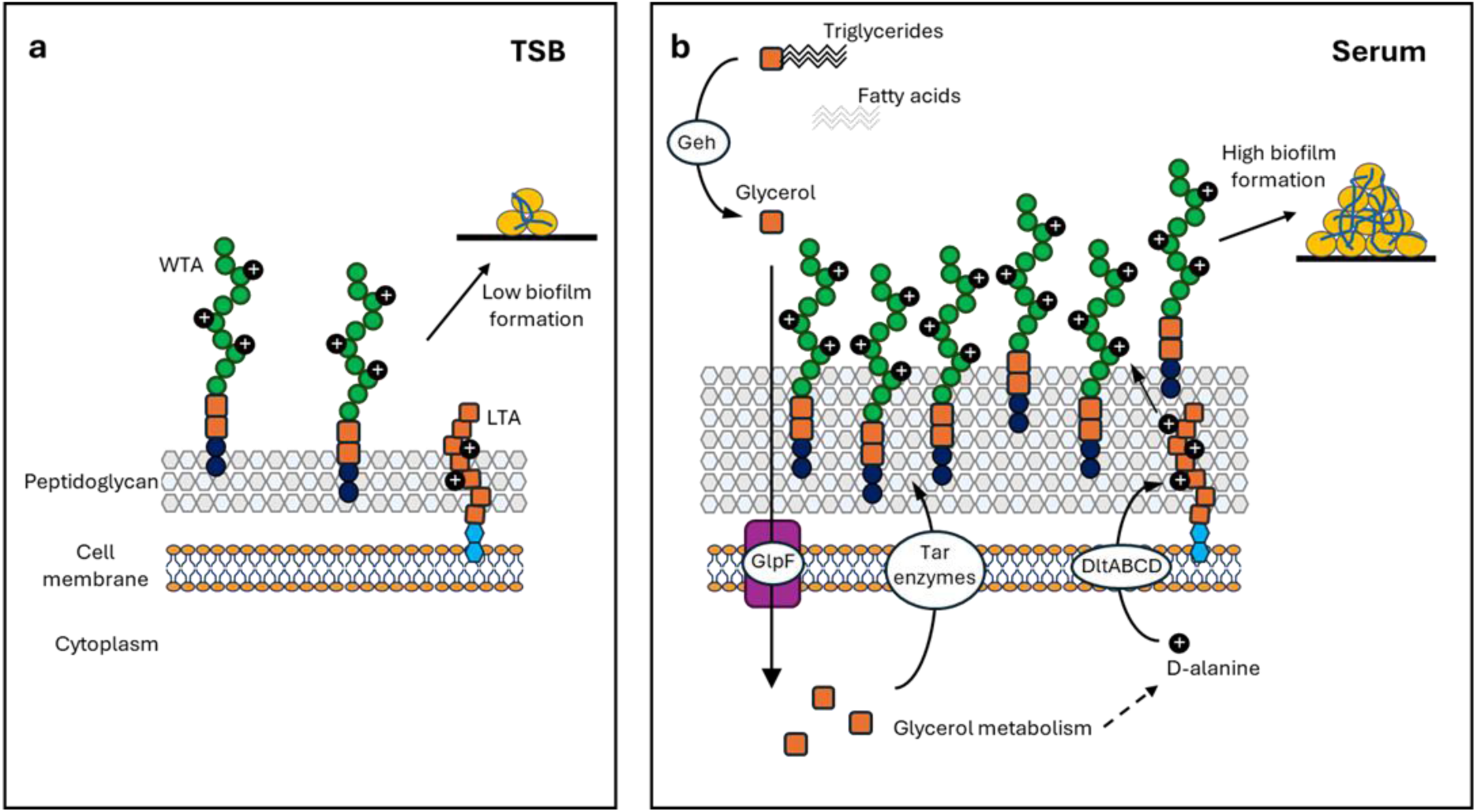
Summary model. (a) In TSB, the staphylococcal peptidoglycan cell wall is decorated with WTA. (b) In human serum, S. *aureus* secretes the Geh lipase which breaks host triglycerides down to release glycerol. After being imported by GlpF and phosphorylated by GlpK, glycerol is further metabolized and then used to support synthesis of increased amounts of WTA, promoting biofilm formation.

There is a growing understanding that the host environment triggers cell wall remodelling in *S. aureus,* including increased peptidoglycan and WTA accumulation^8,9,27^. While the mechanisms underlying the increased peptidoglycan have been defined^8,10^, the basis for increased WTA production was unclear. Here, we show that *S. aureus* uses host lipids to support the enhanced WTA synthesis, demonstrating how host nutrients are not just used for growth but can also be used to alter bacterial physiology and surface properties.

Host-induced cell wall remodelling has profound effects on *S. aureus* behaviour, increasing the antibiotic tolerance of planktonic cells and protecting them from neutrophil-mediated killing^8,11^. Here we show that in addition to this, the increased WTA promotes biofilm formation, further contributing to antibiotic tolerance. This is in line with previous work showing the importance of WTA for adhesion to surfaces and the initial stages of biofilm formation^21,28–30^. WTA are composed of approximately 40 ribitol-5-phosphate molecules covalently attached to the peptidoglycan via a glycerol-containing linkage unit^19^. WTA have been identified as a critical determinant of biofilm formation, although exactly how they mediate this is unknown^21^. They are modified by glycosylation and D-alanylation, the latter of which reduces the net negative charge of the cell surface and affects bacterial surface hydrophobicity. It has been hypothesised that this affects their interaction with surfaces, and in line with this, we observed that D-alanylation was required for the increased biofilm formation observed in serum. As well as direct interactions with surfaces, WTA have also been shown to be required for the secretion of proteins^31^. As under these conditions staphylococcal biofilms are predominantly protein based, increased levels of WTA may lead to increased biofilm biomass by mediating increased release of extracellular proteins. Biofilms have been found to consist of cytoplasmic proteins which associate with the bacterial surface at low pH, however, the mechanism of their release remains unclear^32^.

Our work also uncovers an additional role for the Geh lipase in liberating glycerol to promote cell wall remodelling and biofilm formation. Geh is an important virulence factor, with over 80% of clinical *S. aureus* isolates having lipolytic activity^33^. In addition, lipases are thought to contribute to *S. aureus* dissemination, with isolates from disseminated infections having more lipolytic activity than those from localised infection sites. While lipases have been linked to biofilm formation previously^34^, our work is distinct from this. They have previously been identified to lead to increased biofilm via increasing eDNA release^34^, however, under our conditions, biofilms were susceptible to protease, but not DNase degradation, indicating that the predominant biofilm component was protein and not eDNA. Geh has many other roles as well, including being immunomodulatory by degrading bacterial lipoproteins to hide PAMPs^35^ and detoxifying antibacterial fatty acids^36^.

Using NMR spectroscopy, we directly demonstrate incorporation of the carbon atoms from glycerol into the WTA molecules. This did not just occur in the glycerol moieties present within the linkage unit of WTA but the carbon atoms were also incorporated into the D-alanine and glycosylation modifications, demonstrating extensive metabolism of glycerol and indicating that carbon may an important factor limiting how much WTA *S. aureus* can synthesise. Our data suggest that under these conditions, WTA is predominantly β-glycosylated, with the^13^C NMR spectra showing the presence of one dominant anomeric carbon signal, with a chemical shift more consistent with β-glycosylation than α-glycosylation^24^. This is in line with previous work that found WTA is predominantly α-glycosylated under *in vitro* conditions but β-glycosylated under host conditions^37^. Although we did not find that the type of glycosylation influenced biofilm formation under these conditions, this further highlights the importance of studying *S. aureus* under host-relevant conditions. As the modification of WTA with D-alanine is crucial for the resistance of *S. aureus* to host AMPs and the survival of neutrophil-mediated killing, the finding that glycerol supports this modification raises the possibility that *S. aureus* may exploit host lipids to resist the immune system in addition to influencing biofilm formation.

Our work also links Geh to antibiotic tolerance as the biofilms it led to showed high levels of tolerance to the last resort antibiotic daptomycin. Geh has previously been shown to protect from antibacterials that work by inhibiting fatty acid synthesis for example triclosan, as *S. aureus* can incorporate the released host fatty acids into its own phospholipids^38^. However, this antibiotic tolerance is broader than this as biofilms are typically multi-drug tolerant. Indeed, we observed that the bacteria within serum-formed biofilms showed high levels of tolerance to both vancomycin and daptomycin, antibiotics with distinct mechanisms of action.

This work highlights the importance of studying *S. aureus* under host-relevant conditions. These findings add to the growing body of work demonstrating that the host environment is an active determinant of bacterial physiology^4,8,11^. A failure to account for these host-driven phenotypes may have contributed to an incomplete understanding of biofilm biology and staphylococcal pathogenesis and treatment failure in invasive disease.

Finally, our work has generated a potential viable new combination approach to prevent biofilm formation and to reduce the associated antibiotic tolerance. One approach is to inhibit the lipase enzyme directly. We demonstrated that a loss of the lipase led to a reduction in WTA, reduced biofilm formation and increased susceptibility to daptomycin. This reduced biofilm formation and daptomycin tolerance was recapitulated by inhibiting the lipase enzyme with the clinically used anti-obesity drug orlistat. While orlistat would not be useful in the case of invasive staphylococcal disease due to its inability to reach the bloodstream, this demonstrates the feasibility of this approach. Similarly, we found that inhibition of WTA synthesis using tunicamycin also prevented serum-induced biofilm formation and the associated daptomycin tolerance.

In summary, we demonstrate that adaptation to the host environment actively primes *S. aureus* for enhanced biofilm formation and antibiotic tolerance. We identify the mechanism underlying this, whereby the Geh lipase liberates glycerol from host lipids to promote WTA synthesis and cell wall remodelling. These findings reveal that transit through the bloodstream is not a passive stage of infection but instead alters bacterial surface physiology to promote persistence at secondary sites. By identifying lipase activity and WTA synthesis as critical for this host-induced phenotype, we provide a rationale for therapeutic strategies that target the bacterial pathways specifically engaged in host adaptation, with the potential to prevent biofilm formation *in vivo* and improve the treatment of invasive staphylococcal infections.

## Materials and methods

### Bacterial strains and growth conditions

The bacterial strains used in this study are shown in Table 2. Strains were grown at 37 °C on tryptic soy agar (TSA) or in tryptic soy broth (TSB) with shaking (180 rpm) supplemented with erythromycin (10 µg ml^−1^) or kanamycin (90 µg ml^−1^) when required.

**Table 2.** Strains used in this study.

| Strain | Relevant characteristics | Reference/<br>source |
| --- | --- | --- |
| USA300 JE2 | Community-acquired MRSA strain of the USA300 lineage isolated from a skin and soft tissue infection at the Los Angeles County (LAC) jail, cured of plasmids | 39 |
| SH1000 | *rsbU*<sup>+</sup> derivative of laboratory strain 8325-4 | 40 |
| Col | Early hospital-associated MRSA isolate, commonly used in laboratory experiments | 41 |
| Newman | Clinical isolate, MSSA, used widely in animal and laboratory experiments | 17 |
| ASARM181 | Clinical MRSA endocarditis isolate from CC22 | 13 |
| ASARM142 | Clinical MRSA endocarditis isolate from CC22 | 13 |
| ASASM391 | Clinical MSSA endocarditis isolate from CC30 | 13 |
| ASASM17 | Clinical MSSA endocarditis isolate from CC30 | 13 |
| ASASM351 | Clinical MSSA endocarditis isolate from CC30 | 13 |
| JE2 *geh*::Tn | JE2 defective for Geh lipase. Erm<sup>R</sup> | 39 |
| JE2 *geh*::Tn pCN34 | JE2 defective for Geh lipase complemented with the empty pCN34 vector. Erm<sup>R</sup>; Kan<sup>R</sup> | This study |
| JE2 *geh*::Tn pCN34-*geh* | JE2 defective for Geh lipase complemented with the pCN34 vector containing *geh*. Erm<sup>R</sup>; Kan<sup>R</sup> | This study |
| JE2 *lip3*::Tn | JE2 defective for SAL3. Erm<sup>R</sup> | 39 |
| JE2 *fakA*::Tn | JE2 defective for FakA. Erm<sup>R</sup> | 39 |
| JE2 *fakB1*::Tn | JE2 defective for FakB1. Erm<sup>R</sup> | 39 |
| JE2 *fakB2*::Tn | JE2 defective for FakB2. Erm<sup>R</sup> | 39 |
| JE2 *srtA*::Tn | JE2 defective for SrtA. Erm<sup>R</sup> | 39 |
| JE2 *icaA*::Tn | JE2 defective for IcaA. Erm<sup>R</sup> | 39 |
| JE2 *tarM*::Tn | JE2 defective for TarM. Erm<sup>R</sup> | 39 |
| JE2 *tarS*::Tn | JE2 defective for TarS. Erm<sup>R</sup> | 39 |
| USA300 LAC* | LAC strain of the USA300 CA-MRSA lineage, cured of LAC-p03 plasmid | 42 |
| LAC* Δ*tagO* | LAC* with the *tagO* gene deleted. | 43 |
| LAC* Δ*tagO* pCN34 | LAC* with the *tagO* gene deleted complemented with the empty pCN34 vector. Kan<sup>R</sup> | This study |
| LAC* Δ*tagO* pCN34-*tagO* | LAC* with the *tagO* gene deleted complemented with the pCN34 vector containing *tagO*. Kan<sup>R</sup> | This study |
| LAC* Δ*ltaS*-S2 | LAC*Δ*ltaS*::*erm* suppressor S2. Erm<sup>R</sup>. | 44 |
| LAC* Δ*ltaS*-S2 pCN34 | LAC*Δ*ltaS*::*erm* suppressor S2 complemented with the empty pCN34 vector. Erm<sup>R</sup>; Kan<sup>R</sup> | This study |
| LAC* Δ*ltaS*-S2 pCN34-*ltaS* | LAC*Δ*ltaS*::*erm* suppressor S2 complemented with the pCN34 vector containing *ltaS*. Erm<sup>R</sup>; Kan<sup>R</sup> | This study |
| LAC* Δ*dltD* | LAC* with the *dltD* gene deleted, Erm<sup>R</sup> | 8 |
| LAC* $\Delta dltD$ pCN34 | LAC* with the *dltD* gene deleted, complemented with the empty pCN34 vector. Erm<sup>R</sup>; Kan<sup>R</sup> | 8 |
| LAC* $\Delta dltD$ pCN34-*dltD* | LAC* with the *dltD* gene deleted, complemented with the pCN34 vector containing *dltD*. Erm<sup>R</sup>; Kan<sup>R</sup> | 8 |

To generate TSB-grown bacteria, TSB was inoculated with 10^7^ CFU ml^−1^ from overnight cultures and incubated at 37 °C with shaking (180 rpm) until 10^8^ CFU ml^−1^ was reached. To generate serum-incubated bacteria, these TSB-grown bacteria were centrifuged, resuspended in the same volume of human serum from male AB plasma (Sigma) and incubated for 16 h at 37 °C with shaking (180 rpm). To mitigate differences between batches of serum, each experiment was performed with a single batch of human serum.

As *S. aureus* is unable to replicate in human serum, bacterial CFU counts of serum-incubated cultures were equal to those of TSB-grown cultures (Fig. S1). CFU counts were determined by serial dilution of cultures in PBS and plating onto TSA. Where necessary, serum was supplemented with sub-lethal concentrations of tunicamycin (128 µg ml^−1^) or orlistat (100 µg ml^−1^).

Where appropriate, serum was treated or fractionated before bacterial incubation was carried out. Serum was fractionated via centrifugation through a membrane with a 10 kDa molecular weight cut-off before both the retentate and filtrate were collected and made up to the original volume with PBS. Heat inactivated (HI) serum was generated by heating serum to 56 °C for 30 min. Delipidated serum (DLS) was generated by incubating serum with silica (20 mg ml^-1^, Merck) for 2 h with mixing before centrifugation and filtration of supernatant through a 0.2 µm filter. Where appropriate, DLS was supplemented with lipids extracted from serum using a modified Bligh-Dyer extraction, triolein (1 mM), tripalmitate (1 mM), tributyrin (1 mM), glycerol (0.1%) or fatty acids (5 mM when used individually or 1 mM each fatty acid when used in combination to give a total fatty acid concentration of 5 mM).

### Complementation of the Δ*tagO,* Δ*ltaS-*S2 and *geh*::Tn mutants

Mutants were complemented by inserting the gene amplified from JE2 wildtype along with approximately 500 bp up and downstream of the gene to include the promoter and terminator regions into pCN34. Fragments were amplified using Phusion using the primers shown in Table 3. Fragments were then digested (BamHI/EcoRI for *tagO*, BamHI/SacI for *ltaS* and SalI/SacI for *geh*) and ligated into similarly digested pCN34 using T4 ligase. Ligation mixtures were transformed into *E. coli* IM08B and then electroporated into the relevant mutant along with empty pCN34 plasmid as a control.

**Table 3.**
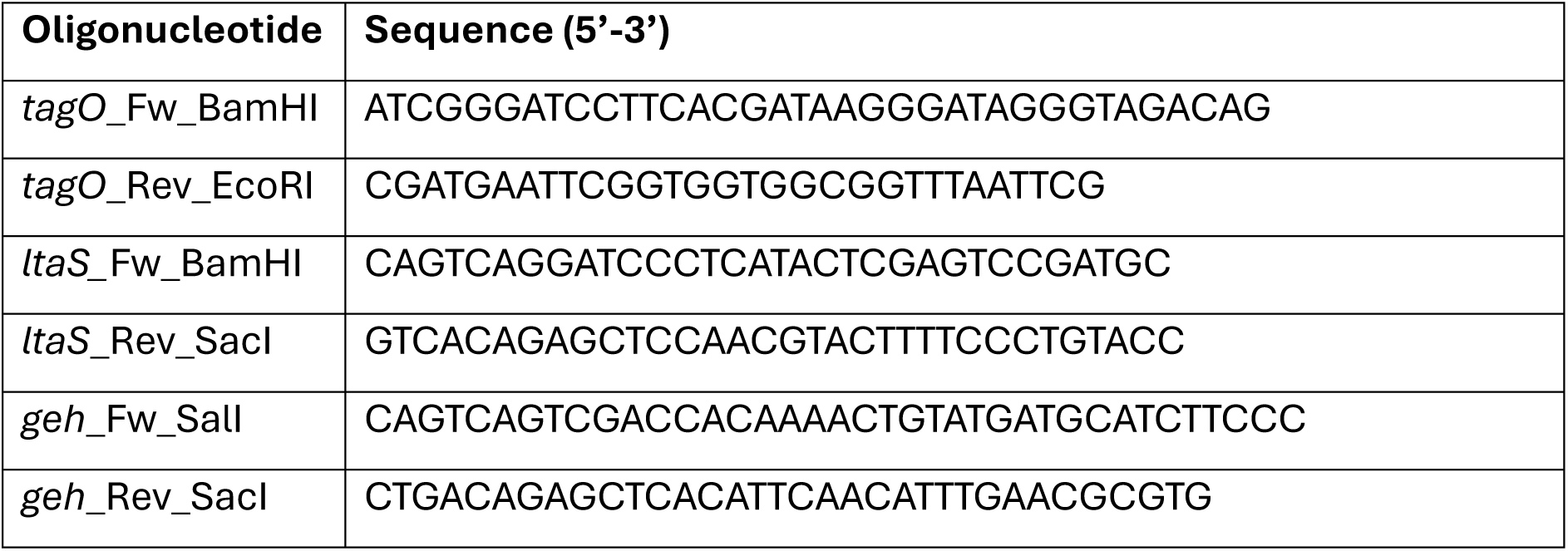
Primers used in this study.

### Biofilm formation and quantification

TSB-grown or serum-incubated *S. aureus* were generated as described above before being washed 3 times in PBS and inoculated at 10^7^ CFU ml^-1^ into individual wells of a flat-bottomed 96 well plate. Each well contained 200 µl TSB supplemented with 1% glucose. Plates were incubated statically for 24 h at 37 °C before biofilm quantification.

Biofilms were washed three times with 300 µl PBS before being incubated with 200 µl 0.1% crystal violet for 15 min statically at room temperature. Unbound crystal violet was removed by three washes with 300 µl PBS. Crystal violet was permeabilised with 150 µl ethanol which was transferred to a new 96 well plate and the absorbance measured at 595 nm using a Tecan M200 PRO plate reader.

To determine the structural components of the biofilm, after biofilm formation as described above, biofilms were washed three times with PBS before being incubated for 3 h at 37 °C with TSB supplemented with 1% glucose and 100 µg ml^-1^ proteinase K, 140 U ml^-1^ DNase or 10 mM sodium metaperiodate. Biofilms were then washed three times in PBS and quantified as above with crystal violet.

### Antibiotic exposure of intact biofilms

Biofilms were generated as described above and washed three times in PBS before 200 µl TSB supplemented with 1% glucose and either 20 µg ml^-1^ daptomycin or 20 µg ml^-1^ vancomycin was added if appropriate. In the case of daptomycin, media were also supplemented with 1.25 mM CaCl_2_. Biofilms were incubated with antibiotics for a further 24 h statically at 37 °C before the metabolic activity was determined using XTT. Biofilms were washed three times in PBS before being incubated with 200 µl PBS containing 0.5 mg ml^-1^ XTT and 20 µM menadione for 30 min statically at 37 °C. Supernatant (150 µl) was moved to a fresh 96 well plate and absorbance determined at 490 nm using a Tecan M200 PRO plate reader.

### Antibiotic tolerance of disrupted biofilms

Biofilms were generated as described above, washed three times in PBS and then resuspended vigorously in 200 µl PBS. This 200 µl was added to 3 ml TSB containing either 20 µg ml^-1^ daptomycin or 20 µg ml^-1^ vancomycin. In the case of daptomycin, media were also supplemented with 1.25 mM CaCl_2_. Samples were incubated at 37 °C with shaking (180 rpm) for 6 h. At each time-point (0, 2, 4, 6 h), aliquots were taken, serially diluted 10-fold in PBS and plated onto TSA for CFU counts.

### WTA extraction and quantification

Serum-incubated bacteria were generated as described above and WTA was extracted as described previously. 5 ml cultures were washed with 10 ml 50 mM MES (pH 6.5) (Buffer 1) and resuspended in 15 ml 50 mM MES (pH 6.5) supplemented with 4% SDS (Buffer 2).

Samples were boiled for 1 h and centrifuged before being washed twice in 1 ml Buffer 2, once in 1 ml 50 mM MES (pH 6.5) supplemented with 2% NaCl (Buffer 3), and once in 1 ml Buffer 1. The pellet was resuspended in 1 ml 20 mM Tris-HCl pH 8, 0.5% SDS and digested with 20 µg proteinase K for 4 h at 50 °C. The pellet was washed once with 1 ml Buffer 3, three times with 1 ml water, resuspended in 250 µl 0.1 M NaOH and incubated for 16 h at room temperature. After centrifugation, supernatant was neutralised with 62.5 µl 1 M Tris-HCl (pH 7.8) and analysed by PAGE. 20 µl aliquots of WTA samples were separated on a 20% native polyacrylamide gel by electrophoresis using 0.1 M Tris, 0.1 M Tricine, pH 8.2 running buffer. Gels were then stained with alcian blue (1 mg ml^-1^, 3% acetic acid), destained with water and quantified using ImageJ.

### NMR spectroscopy

NMR spectra were recorded at 150.9 MHz on a Bruker AVIII 600 NMR spectrometer equipped with a 5 mm BB H&F cryoprobe using a ^13^C uniform driven equilibrium Fourier transform (UDEFT) pulse sequence for increased detection of quaternary carbon signals^45^. The spectra were acquired at 25 °C with 8192 scans and 65536 data points, using a relaxation delay of 5 s and a spectral width of 240 ppm. Prior to acquisition, 10% v/v D_2_O was added to the extracted WTA samples for the purpose of locking and shimming. Spectra were processed using Topspin 3.7 software and signals were referenced to the chemical shift of the tris *C*H_2_ signal at 61.4 ppm^25^.

### Statistical analyses

The CFU data were log_10_ transformed. The data were analysed via a Student’s *t* test, one-way analysis of variance (ANOVA), or two-way ANOVA with a *post hoc* test to correct for multiple comparisons, as is described in the figure legends, using GraphPad Prism (v10.0).

## Acknowledgements

All authors acknowledge the provision of strains by the Network on Antimicrobial Resistance in *Staphylococcus aureus* (NARSA) Program: under NIAID/NIH Contract number HHSN272200700055C. This work was supported by Research Ireland Frontier for the Future Program Award (reference: 21/FFP-A/9704), a Wellcome Trust Investigator Award (reference: 212258/Z/18/Z) and a Microbiology Society Vacation Studentship (reference: GA004851).

