## Supplementary data for "Human serum triglycerides promote *Staphylococcus aureus* biofilm formation and antibiotic tolerance"

**Supplementary figures 1-14**

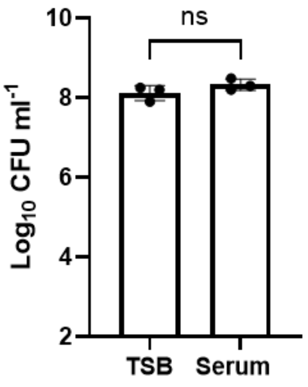


**Fig. S1. Bacterial numbers do not increase during incubation in serum.** There was no significant difference in bacterial survival as measured by $log_{10}$ CFU $ml^{-1}$ of the JE2 WT strain after growth in TSB and after the 16 h incubation in serum. Data represent the geometric mean ± geometric standard deviation of three independent repeats and were analysed by t-test (ns, $p > 0.05$).

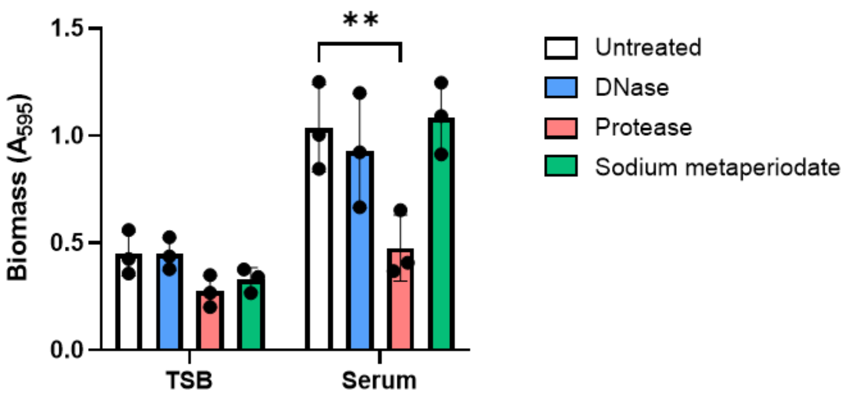


**Fig. S2. Biofilms are predominantly composed of proteins.** Biofilms were formed from TSB-grown and serum-incubated JE2 WT for 24 h, then incubated for a further 3 h with 140 U $ml^{-1}$ DNase, 100 µg $ml^{-1}$ proteinase K or 10 mM sodium metaperiodate to degrade DNA, protein and carbohydrates, respectively, before quantification of remaining biomass with crystal violet. Data represent the mean ± standard deviation of three independent repeats and were analysed by two-way ANOVA with Dunnett's *post-hoc* test. **, $P < 0.01$. For all other tested comparisons (untreated vs treated), $P > 0.05$.

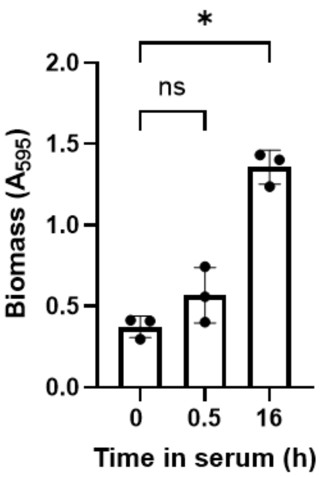


**Fig. S3. Increased biofilm formation in serum is not fully explained by host factors sticking to *S. aureus*.** JE2 WT was grown in TSB and then incubated in serum for 0, 0.5 or 16 h before being inoculated into TSB and incubated for 24 h to form biofilm before quantification of biofilm biomass with crystal violet. Data represent the mean ± standard deviation of three independent repeats and were analysed by one-way ANOVA with Sidak's *post-hoc* test. ns, $P > 0.05$, *, $P < 0.05$.

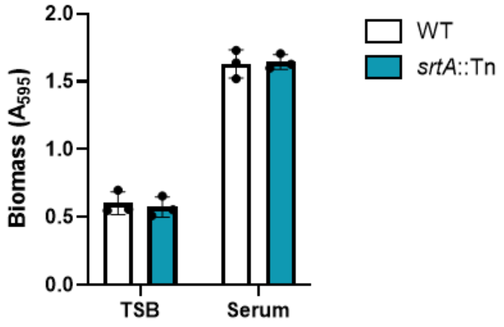


**Fig. S4. Cell wall anchored proteins are not required for biofilm formation in serum.** The JE2 WT strain and the *srtA*::Tn mutant were TSB-grown and serum-incubated before being inoculated into TSB and incubated for 24 h to form biofilm, which was then quantified using crystal violet. Data represent the mean ± standard deviation of three independent repeats and were analysed by two-way ANOVA with Sidak's *post-hoc* test, for all WT vs mutant comparisons, $P > 0.05$.

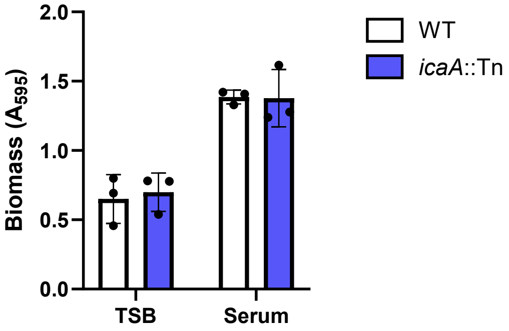


**Fig. S5. PIA is not required for biofilm formation in serum.** The JE2 WT strain and the *icaA*::Tn mutant were TSB-grown and serum-incubated before being inoculated into TSB and incubated for 24 h to form biofilm, which was then quantified using crystal violet. Data represent the mean ± standard deviation of three independent repeats and were analysed by two-way ANOVA with Sidak's *post-hoc* test, for all WT vs mutant comparisons, $P > 0.05$.

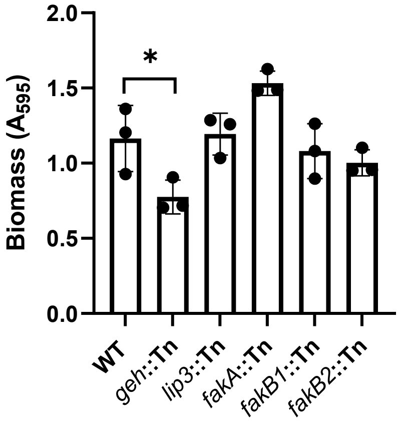


**Fig. S6. Geh but not fatty acid kinase is required for biofilm formation in serum.** The JE2 WT strain and the *geh*::Tn, *lip3*::Tn, *fakA*::Tn, *fakB1*::Tn and *fakB2*::Tn mutants were serum-incubated before being inoculated into TSB and incubated for 24 h to form biofilm, which was then quantified using crystal violet. Data represent the mean ± standard deviation of three independent repeats and were analysed by one-way ANOVA with Dunnett's *post-hoc* test. *, $P < 0.05$.

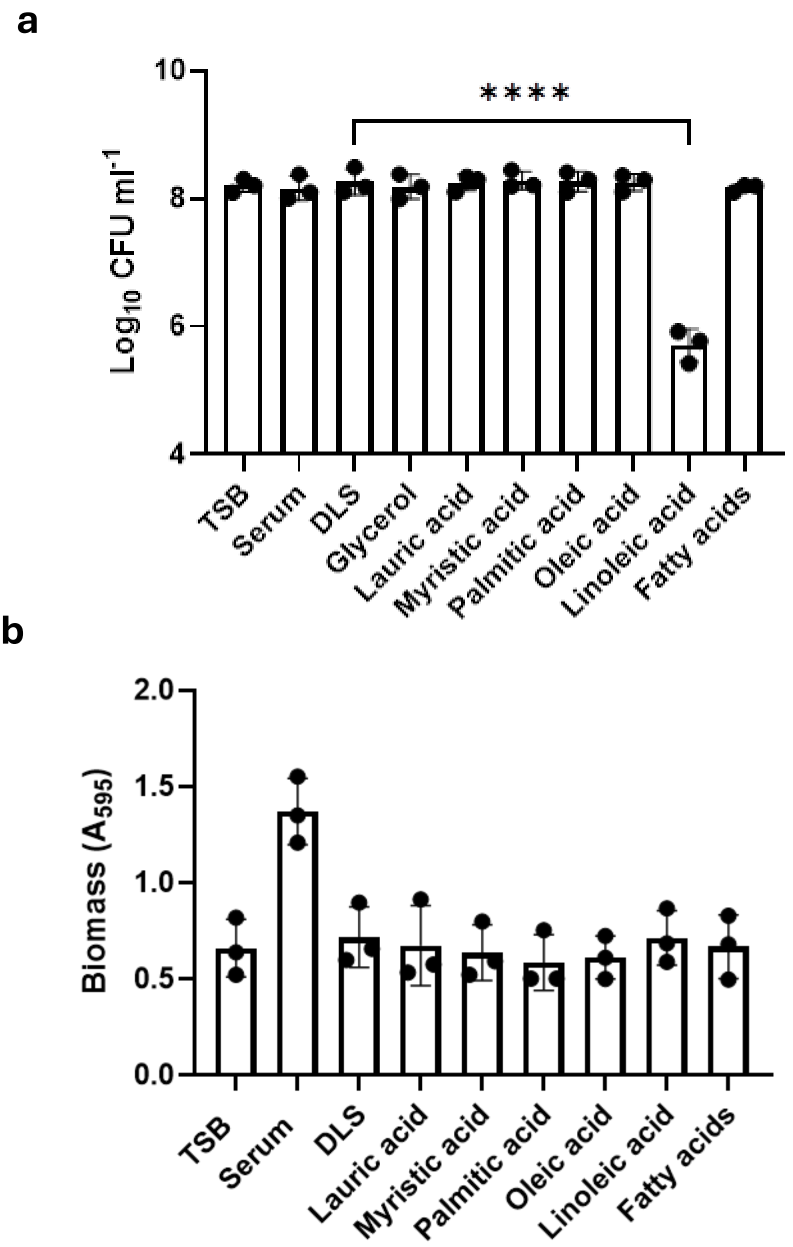


**Fig. S7. Addition of fatty acids does not promote biofilm formation.** JE2 WT was TSB-grown, serum-incubated or incubated in DLS supplemented or not with 1 mM individual fatty acids or 5 mM total fatty acids before (a) $\log_{10}$ CFU $ml^{-1}$ were determined and (b) bacteria were inoculated into TSB, incubated for 24 h to form biofilm and biofilm quantified with crystal violet. Data represent the mean ± standard deviation of three independent repeats. Data were analysed by one-way ANOVA with Dunnett's *post-hoc* test. In panel a, there were no significant differences between bacterial survival in DLS and any other condition except linoleic acid (*, $P < 0.05$). In panel b, there was no significant difference between biomass in DLS or DLS supplemented with any fatty acid.

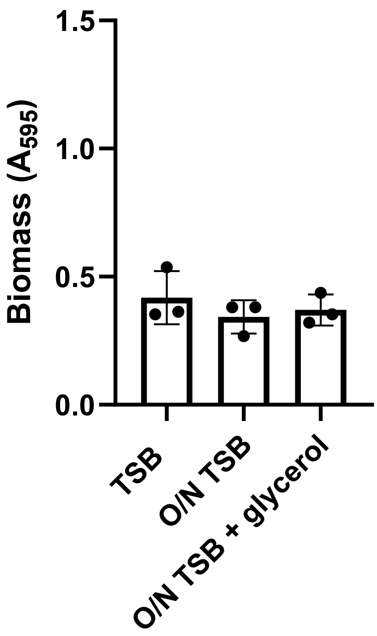


**Fig. S8. Addition of glycerol into TSB does not promote biofilm formation.** Biofilm biomass of JE2 WT grown to mid-exponential phase in TSB, grown for 16 h in TSB or for 16 h in TSB supplemented with 0.1% glycerol. Data represent the mean ± standard deviation of three independent repeats. There was no significant difference in biofilm biomass observed under any condition (data were analysed by one-way ANOVA with Tukey's *post-hoc* test, for all comparisons, $P > 0.05$).

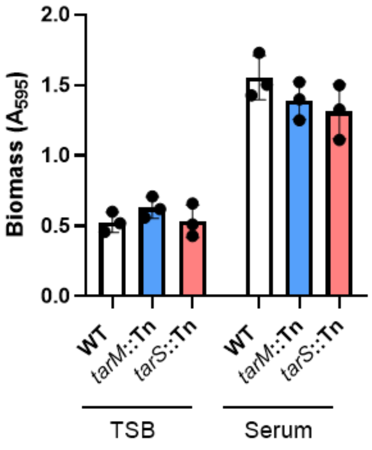


**Fig. S9. WTA glycosylation is not required for biofilm formation.** The JE2 WT strain and the *tarM*::Tn and *tarS*::Tn mutants were TSB-grown and serum-incubated before being inoculated into TSB and incubated for 24 h to form biofilm, which was then quantified using crystal violet. Data represent the mean ± standard deviation of three independent repeats and were analysed by two-way ANOVA with Sidak's *post-hoc* test. For all comparisons (WT vs mutants under each condition), $P > 0.05$.

**Fig. S10. Glycerol supplementation promotes WTA synthesis in delipidated serum.**

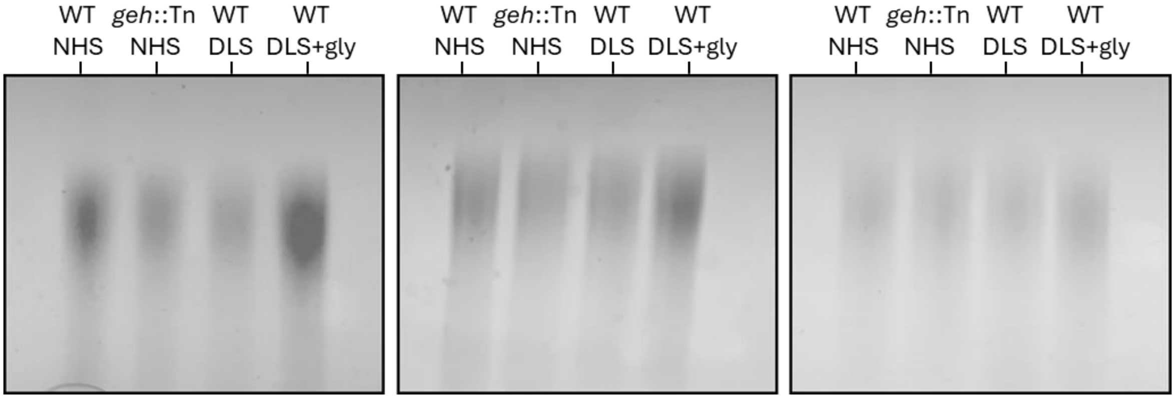


Native PAGE analysis of WTA extracted from JE2 WT and the *geh*::Tn mutant after incubation in NHS, DLS or DLS supplemented with glycerol. These gels were quantified using ImageJ to generate the graph in Fig. 3f. NHS, normal human serum; DLS, delipidated serum; gly, glycerol.

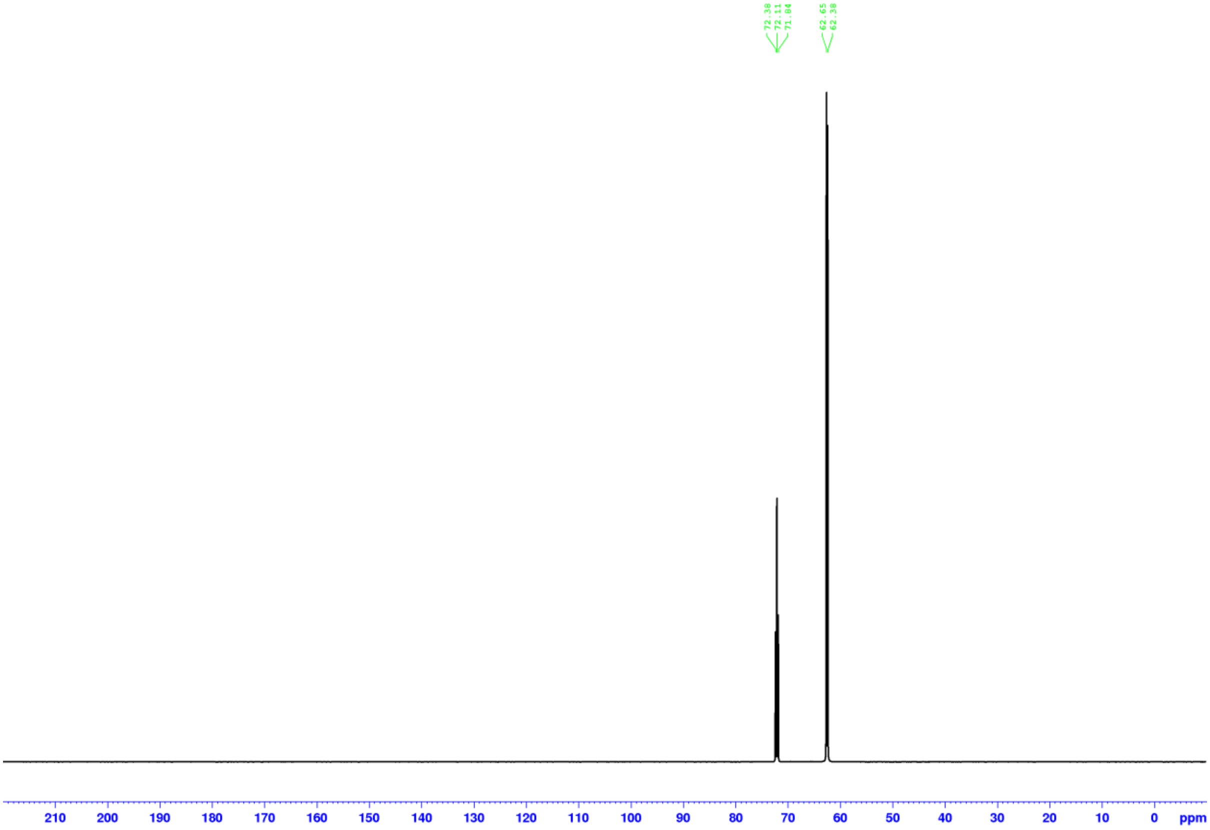


**Fig. S11.** $^{13}$**C NMR (UDEFT, 90:10** $H_2O/D_2O$**, 150.9 MHz) spectrum of** $^{13}$**C-glycerol.**

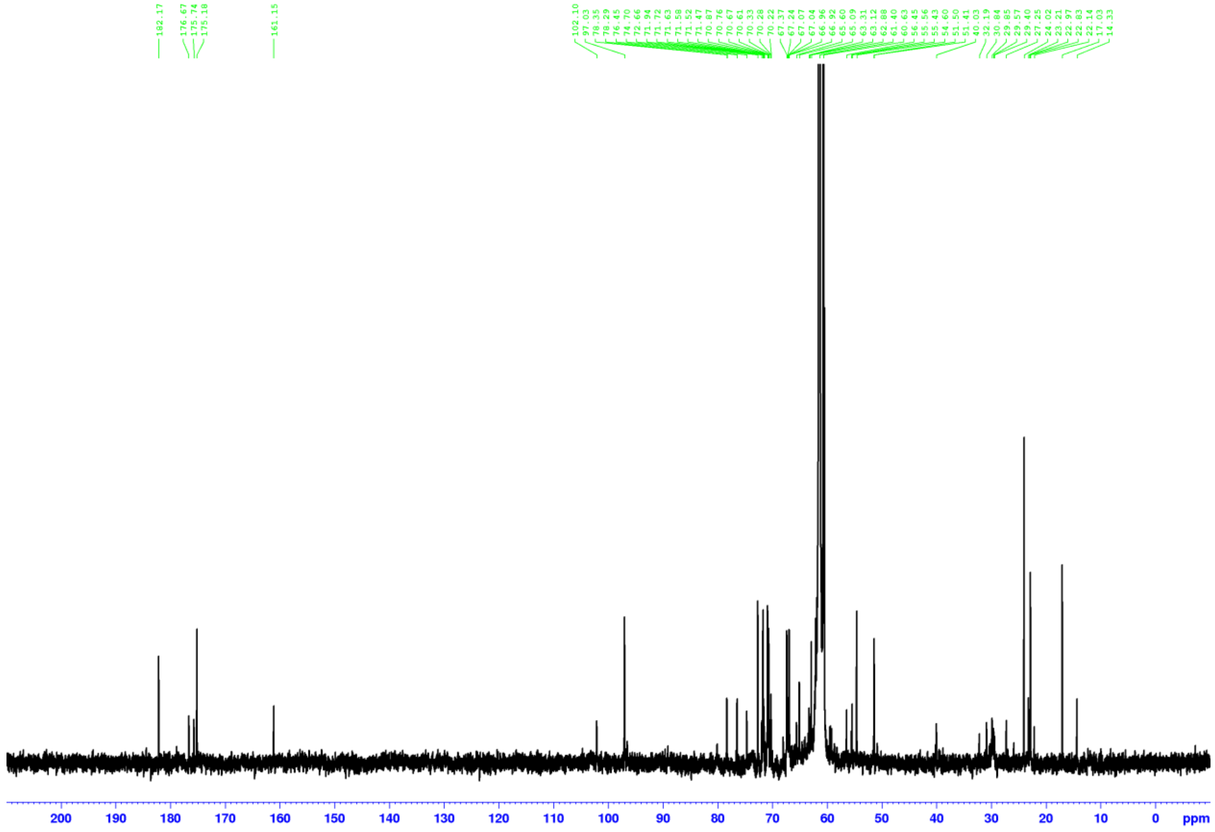


**Fig. S12. $^{13}C$ NMR (UDEFT, 90:10 $H_2O/D_2O$, 150.9 MHz) spectrum of DLS + glycerol sample (WTA extraction in 0.2 M tris HCl).**

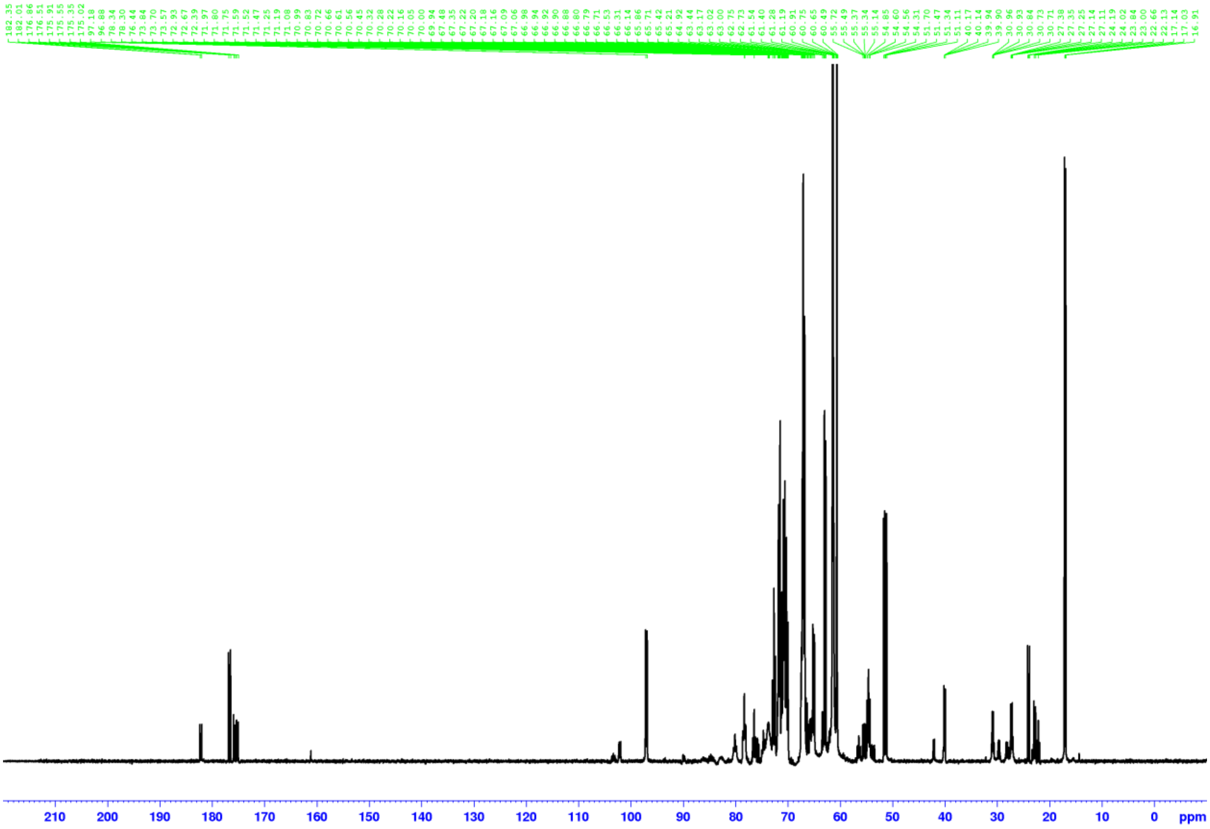


**Fig. S13.** $^{13}C$ **NMR (UDEFT, 90:10** $H_2O/D_2O$**, 150.9 MHz) spectrum of DLS +** $^{13}C$**-glycerol sample (WTA extraction in 0.2 M tris HCl).**

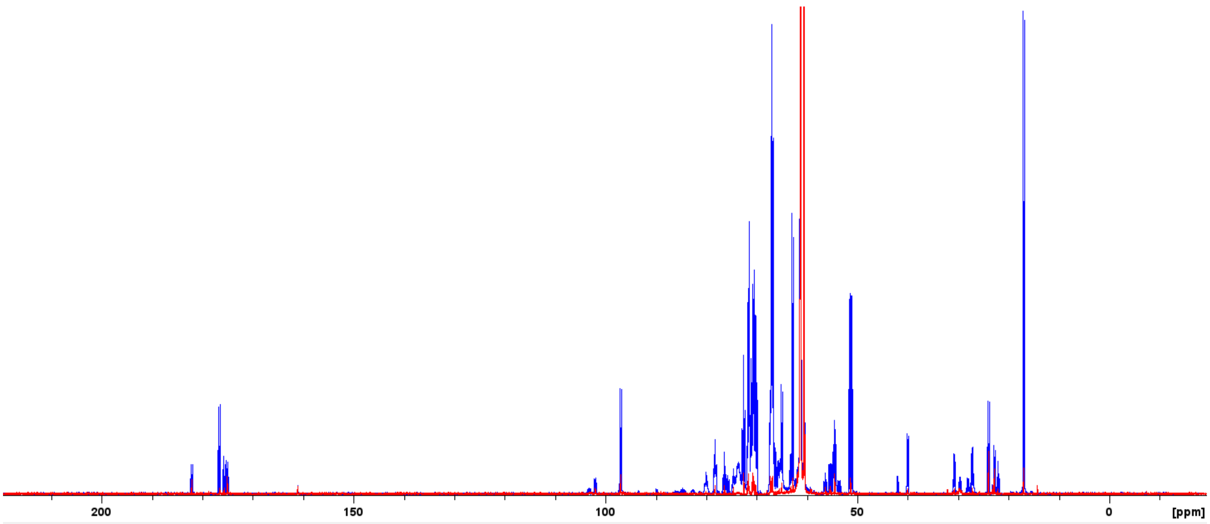


**Fig. S14. Directly overlain $^{13}$C NMR (UDEFT, 90:10 $H_2O/D_2O$, 150.9 MHz) spectra of DLS + $^{13}$C-glycerol sample (WTA extraction in 0.2 M tris HCl) (blue) and DLS + C-glycerol sample (WTA extraction in 0.2 M tris HCl) (red).** The tris signals are scaled to the same intensity for both spectra shown.
